# The effect of a lifestyle intervention in obese pregnant women on change in gestational metabolic profiles: findings form the UK Pregnancies Better Eating and Activity Trial (UPBEAT) RCT

**DOI:** 10.1101/125740

**Authors:** Harriet Mills, Nashita Patel, Sara White, Dharmintra Pasupathy, Annette L Briley, Diana dos Santos Ferreira, Paul T Seed CStat, Scott M. Nelson, Naveed Sattar, Kate Tilling, Lucilla Poston, Debbie A Lawlor, On behalf of the UPBEAT Consortium.

**Author notes:** **Corresponding Author** DA Lawlor, MRC Integrative Epidemiology Unit at the University of Bristol, Oakfield House, Oakfield Grove, Bristol, BS8 2BN, UK.

## Abstract

**Background:** Pregnancy metabolic disruption is believed to be enhanced in obese women and lead to adverse outcomes in them and their offspring. The UK Pregnancies Better Eating and Activity Trial (UPBEAT), a randomised controlled trial of a lifestyle intervention in obese pregnant women, has already been shown to improve diet and physical activity. We used UPBEAT to determine (a) the magnitude of change in metabolic profiles in obese pregnant women and (b) the impact of a lifestyle intervention that successfully improved diet and physical activity on these profiles.

**Methods:** Detailed targeted metabolic profiling, with quantification of 158 metabolic features (129 lipid measures, 9 glycerides and phospholipids, and 20 low-molecular weight metabolites) was completed on three occasions (∼17-, 28- and 35-weeks of gestation) in the UPBEAT participants using NMR. Random intercept and random slope models were used to quantify metabolite changes in obese women using the control (usual care) group only (N = 577). The effect of the intervention was determined by comparing rates of metabolite change between those randomised to intervention and usual care, using intention to treat analyses (N = 1158).

**Results:** There were adverse changes across pregnancy in most lipoprotein subclasses, lipids, glycerides, phospholipids, several fatty acids and glucose. All extremely large, very large, large, medium, small and very small VLDL particles increased by 2 to 3 standard deviation units (SD), with IDL, and large, medium and small LDL particles increasing by 1 to 2SD, between 16- and 36-weeks. Triglycerides increased by 3 to 4SD and glucose increased by 2SD, with more modest changes in other metabolites. The intervention reduced the rate of increase in extremely large, very large, large and medium VLDL, in particular those containing triglycerides. Triglyceride to phosphoglyceride ratio was reduced and there were improvements in fatty acid profiles (increases in the proportion of all fatty acids that were linoleic, omega-6 and polyunsaturated and decreases in the proportion of saturated).

**Conclusion:** Systemic metabolism is markedly disrupted in obese pregnant women, but a lifestyle intervention that improved their diet and physical activity has beneficial effects on some of these profiles; these effects might have long-term benefit.

## Introduction

Normal pregnancy is associated with marked changes in maternal metabolism that are necessary for healthy fetal growth and development, but which might cause adverse pregnancy, perinatal and longer-term maternal and offspring outcomes if these changes are extreme.^1-4^ It has been suggested that the metabolic changes seen in normal pregnancy are exacerbated in women who are obese and that this increased metabolic disruption mediates at least some of the adverse short- and long-term outcomes associated with obese pregnancy.^1,5-7^ Randomised controlled trials (RCTs) in pregnant women have suggested some beneficial effect of lifestyle interventions on maternal pregnancy adiposity,^8,9^ but whether this improves their metabolic profiles is unclear. A recent RCT of a diet and physical activity lifestyle intervention in 376 obese (BMI > 30kg/m^2^) pregnant women found beneficial effects on gestational weight gain and C-reactive protein, but no evidence of benefit on insulin sensitivity/glucose tolerance or standard lipid (total cholesterol, VLDLc, LDLc, HDLc or triglycerides) measurements.^10^ Biomarkers were only assessed at two time points and analyses did not look at longitudinal change but treated results for each time-point as independent outcomes.

The UK Pregnancies Better Eating and Activity Trial (UPBEAT) RCT tested the effect of an intense behaviour change intervention in 1554 obese pregnant women (BMI > 30kg/m^2^) on adverse pregnancy and perinatal outcomes.^9^ The intervention was effective in reducing dietary intake of total energy, total fat, saturated fat and carbohydrate, as well as achieving a diet with a lower glycaemic load and index, and increasing protein and fibre intake.^9^ It also led to increased amounts of time spent walking and in the metabolic equivalent ratio of activity to rest, but did not have an effect on the primary outcomes of gestational diabetes (GDM) or large for gestational age (LGA).^9^ The recent addition of repeat (three occasions) gestational metabolic measurements in participants from this RCT provides a unique opportunity to determine the extent to which metabolic profiles change across pregnancy in obese women and whether an intervention with known beneficial effects on diet, physical activity and adiposity influences these changes.

The aims of this study were to determine: (a) how metabolic profiles change over gestation in obese pregnant women, and (b) the effect of an intervention that resulted in healthy diet and physical activity changes in obese pregnant women on gestational change in metabolic profiles.

## Methods

### Study design, randomisation and participants

UPBEAT was a multicentre, RCT of a complex behavioural intervention of diet and physical activity advice versus standard antenatal care in obese pregnant women to prevent GDM and delivery of LGA.^9^ It involved eight urban UK centres; in London (three centres), Bradford, Glasgow, Manchester, Newcastle, and Sunderland. Approvals were obtained from the UK research ethics committee (UK integrated research application system, reference 09/H0802/5) and local Research and Development (R and D) departments in participating centres; all women provided written informed consent prior to entering the study. UPBEAT is registered with Current Controlled Trials, ISRCTN89971375.

UPBEAT recruited and randomised 1554 obese women (BMI ≥30 kg/m^2^), aged 16 years or older, and with a singleton pregnancy between 15^+0^ and 18^+6^ weeks of gestation (hereafter weeks). Exclusion criteria were multiple pregnancy, current use of metformin, unwilling or unable to provide written informed consent, or underlying disorders.^9^ Women were randomised using an internet-based, computer generated sequence that ensured concealment. In order to reduce differences between groups occurring by chance the randomisation procedure included minimising by age, ethnic origin, centre, BMI, and parity.^9^ During pregnancy, women were followed up at 27^+0^ to 28^+6^ weeks (when an oral glucose tolerance test (OGTT) was completed) and at 34^+0^ to 35^+6^ weeks.

For the purposes of this study, women from two centres (King’s College Community and Sunderland) were excluded as no blood samples were taken from participants in these centres for resource and logistical reasons (n = 360 in total; 178 and 182, respectively from control and intervention arm were excluded). From the remaining 6 centres all women with at least one metabolic profile assessment were included. Of the 1194 from the remaining 6 centres 1158 (97%) were included, with similar proportions from the intervention and control arm of the trial (**Figure 1**).

**Figure 1:**
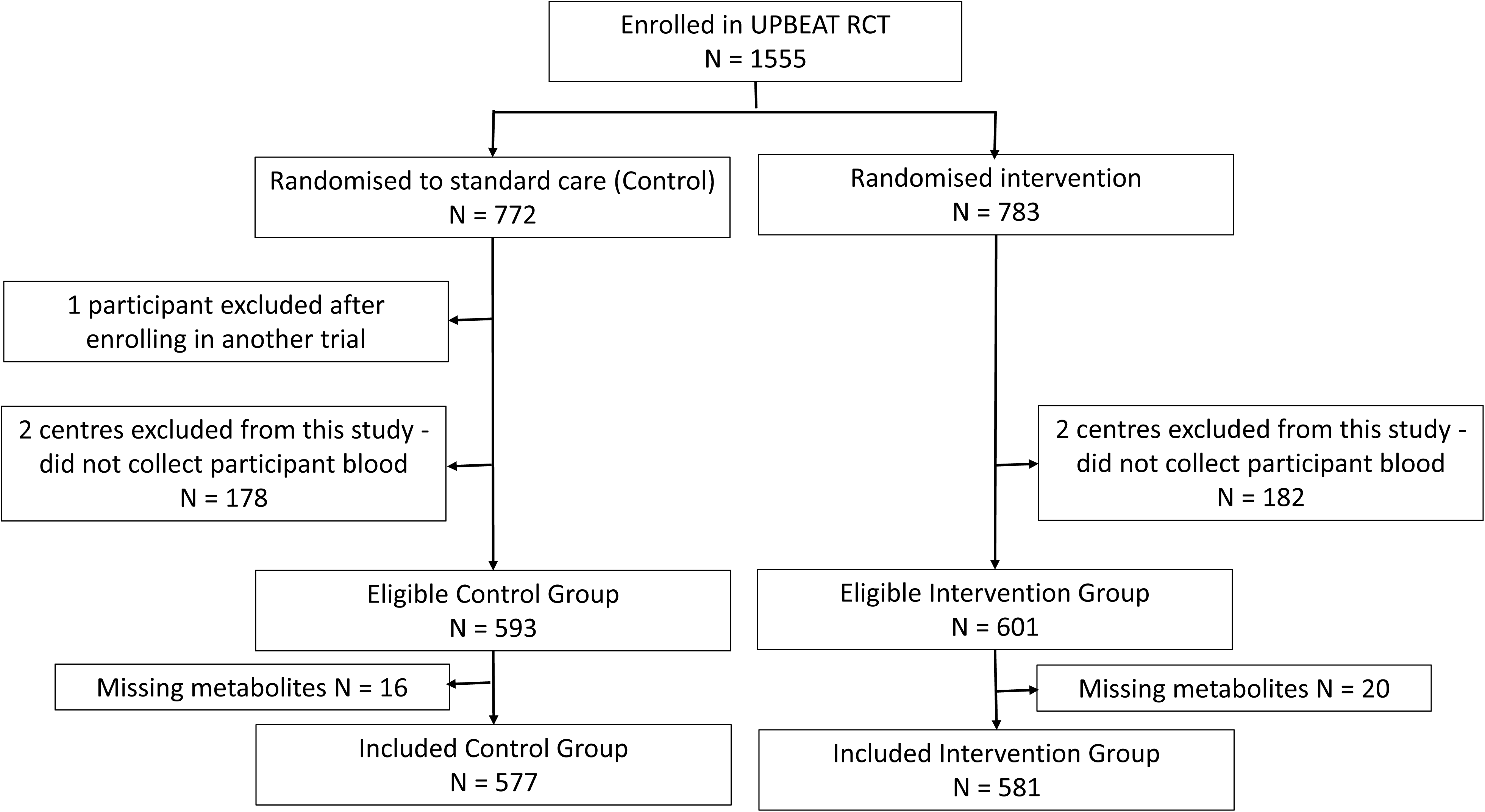
Participant flow.

### Metabolic profiling

Venous blood samples were taken on three occasions: at recruitment, prior to randomization (median (IQR) gestational age: 17.0 (16.1, 17.9) weeks); post-randomisation at the time of the OGTT (27.7 (27.3, 28.1)) and during the third trimester (34.7 (34.3, 35.1)). Samples taken prior to randomization and in the third trimester were non-fasting; those taken as part of the OGTT were after an overnight fast. All blood samples were initially kept on dried ice, processed within 2-hours and then stored at -80oC until used for metabolic profiling. 158 metabolic features were measured and quantified using an NMR targeted platform (http://www.computationalmedicine.fi/), including 129 lipid measures (lipoprotein particle subclasses, particle size, cholesterols, fatty acids and apolipoproteins), 9 glycerides and phospholipids, and 20 low-molecular weight metabolites including branched chain and aromatic amino acids, glycolysis-related markers, and ketone bodies.^11^ This platform has been used in several large scale epidemiological studies,^4,12-15^ and further details of it and a full list of the 158 metabolites are presented in online supplementary material (**sText; sTable 2; sFigure 1**). All blood samples were processed by laboratory technicians blinded to participant data, including which arm of the trial they had been randomised to.

### Statistical analyses

Full details of the statistical modelling and code used to generate results are provided in online supplementary material (**sText**).

Multilevel (random intercept and random slope) models were used to analyse the repeatedly assessed metabolic traits.^16,17^ We restricted the time frame of these models to 16- to 36-weeks so that we were not predicting beyond available data. These models provide an individual (predicted) level of each metabolite at 16-weeks (the intercept) and an individual slope, which we present as the change in each metabolite per 4-weeks increase in gestational age between 16- and 36-weeks. In all analyses we controlled for the minimising variables used in randomization (BMI, ethnicity, parity, age and clinic centre).^9^ An interaction term between time (gestational weeks) and randomised arm (control or intervention) was also included.

### Change in metabolic profiles across pregnancy in obese women in the control arm

We estimated the absolute mean difference in each metabolite between 16- and 36-weeks by subtracting the predicted (from the multi-level model) value at 16-weeks from that at 36-weeks for each women in the control group. These results are presented in standard deviation (SD) units, to aid comparison of results with those from other studies. In these analyses the magnitude of the SD for each metabolite was that from levels at 16-weeks. Thus, if the SD for one metabolite at 16 weeks is 0.5mmol/l, we divided levels at 36-weeks for that metabolite by 0.5mmol/l. We also present the mean absolute differences in the original units of measure for each metabolite (mostly mmol/l).

The full model results (mean intercept and mean slope per 4-weeks) for each metabolite in their original units are also presented for all 1158 women included in analyses. As the model includes a term for the randomised arm for each woman, these can be interpreted as the mean level of each metabolite at 16-weeks, and its change per 4-weeks of gestation between 16- and 36-weeks having adjusted for any effect of the intervention. The slope is therefore an indication of mean rate of change in metabolites in obese women in general (i.e. without any intervention effect).

### Effect of the intervention on change in metabolic profiles across pregnancy

In the multi-level model described above the interaction term between intervention group and gestation weeks represents the mean difference in the rate of change (slope) for each metabolic measure between 16- and 36-weeks between women in the intervention and control groups. It therefore provides the effect of the intervention on rate of change in metabolites. In the main analyses we express this in SD units per 4-weeks (using the SD value of the control group), for ease of interpretation and to enable effects of the intervention to be compared across different metabolic measures. We also present the difference in mean rate of change in original units (mostly mmol/l per 4-weeks).

### Sensitivity analyses

Our main analyses assumes that effect of the intervention is consistent between the first two measurements (∼16-28 weeks) and the second two (∼28 to 36 weeks). To test this assumption we modified the multilevel model to include a knot point at 28-weeks and compared the effect of the intervention for each trait between 16 to 28 week and 28 to 36 weeks.

## Results

Participant characteristics were similar between control and intervention arms and also between those included in this study and those from the six centres where blood samples had been taken and who were eligible to be included in this study (**Table 1**); they were also similar when compared to all women who were randomised irrespective of whether or not they were from a centre where blood sampling was undertaken (**sTable 1**). All women with at least one metabolic profile measure we included in this study. The proportion of those with a metabolic profile measure decreased over gestation in line with a small loss to follow-up in the main RCT, but this was similar in intervention and control groups (**Table 1**).

**Table 1:** Participant characteristics.

|  | **Participants with at least one metabolic profile analysed during pregnancy (analysis sample for this study).**<br>**N = 1158<sup>a</sup>** |  | **All participants in the six centres with blood sampling (eligible sample for this study).**<br>**N = 1194<sup>a</sup>** |  |
| --- | --- | --- | --- | --- |
|  | **Control**<br>**N = 577** | **Intervention**<br>**N = 581** | **Control**<br>**N = 593** | **Intervention**<br>**N = 601** |
| **BMI (N (%))** |  |  |  |  |
| 30 to 34.9 kg/m<sup>2</sup> | 273 (47.3) | 287 (49.4) | 279 (47) | 296 (49.3) |
| 35 to 39.9 kg/m<sup>2</sup> | 203 (35.2) | 177 (30.5) | 209 (35.2) | 185 (30.8) |
| ≥40 kg/m<sup>2</sup> | 101 (17.5) | 117 (20.1) | 105 (17.7) | 120 (20) |
| **Ethnicity (N (%))** |  |  |  |  |
| White | 389 (67.4) | 384 (66.1) | 396 (66.8) | 397 (66.1) |
| Asian | 38 (6.6) | 43 (7.4) | 43 (7.3) | 45 (7.5) |
| Black | 120 (20.8) | 127 (21.9) | 123 (20.7) | 130 (21.6) |
| Other | 30 (5.2) | 27 (4.6) | 31 (5.2) | 29 (4.8) |
| **Parity (N (%))** |  |  |  |  |
| Primiparous | 260 (45.1) | 257 (44.2) | 266 (44.9) | 265 (44.1) |
| Multiparous | 317 (54.9) | 324 (55.8) | 327 (55.1) | 336 (55.9) |
| **Age (N (%))** |  |  |  |  |
| <25 years | 97 (16.8) | 85 (14.6) | 100 (16.9) | 90 (15) |
| 25 to 29 years | 141 (24.4) | 165 (28.4) | 147 (24.8) | 169 (28.1) |
| 30 to 34 years | 187 (32.4) | 174 (29.9) | 188 (31.7) | 182 (30.3) |
| ≥35 years | 152 (26.3) | 157 (27) | 158 (26.6) | 160 (26.6) |
| **Centre (N (%))** |  |  |  |  |
| Bradford | 19 (3.3) | 22 (3.8) | 25 (4.2) | 28 (4.7) |
| Glasgow | 130 (22.5) | 132 (22.7) | 131 (22.1) | 134 (22.3) |
| Manchester | 67 (11.6) | 67 (11.5) | 70 (11.8) | 69 (11.5) |
| Newcastle | 120 (20.8) | 116 (20) | 122 (20.6) | 120 (20) |
| St George's, London | 53 (9.2) | 55 (9.5) | 54 (9.1) | 57 (9.5) |
| St Thomas's, London | 188 (32.6) | 189 (32.5) | 191 (32.2) | 193 (32.1) |
| **First clinic** |  |  |  |  |
| N (%) | 538 (93.2) | 545 (93.8) | 593 (100.0) | 601 (100.0) |
| Median (IQR)<br>gestation (weeks) | 17.0<br>(16.1, 17.9) | 17.0<br>(16.1, 18.0) | 17.0<br>(16.1, 17.9) | 17.0<br>(16.1, 18.0) |
| **Second clinic** |  |  |  |  |
| N (%) | 500 (86.7) | 477 (82.1) | 591 (99.7) | 598 (99.5) |
| Median (IQR)<br>gestation (weeks) | 27.7 (27.3,<br>28.1) | 27.7<br>(27.3, 28.1) | 27.7<br>(27.3, 28.3) | 27.7<br>(27.3, 28.1) |
| **Third clinic** |  |  |  |  |
| N (%) | 407 (70.5) | 374 (64.4) | 524 (88.4) | 485 (80.7) |
| Median (IQR)<br>gestation (weeks) | 34.7<br>(34.3, 35.1) | 34.6<br>(34.3, 35.1) | 34.7<br>(34.3, 35.3) | 34.7<br>(34.3, 35.3) |

<sup>a</sup> The 1158 participants whose results are in the first two columns are a subgroup of the 1194 whose results are presented in the last two columns. N: number; IQR: Interquartile range

### Change in metabolic profiles in obese pregnant women

Absolute concentrations of all lipids, phospholipids and triglycerides in lipoprotein subclasses, with the exception of large, medium and small HDL, increased across pregnancy from 16- to 36-weeks in the control women (**sTable 3**). Most of these changes were substantial, with lipids and phospholipids in all sizes of VLDL, IDL and LDL concentrations, increasing by 1-3SDs, and concentrations of triglycerides in these lipoproteins increasing by 3-4SDs. Concentrations of very large HDL particles increased between 0.3 to 0.5SD, except for triglycerides in these particles which increased by ∼3SD. Concentrations of most large, medium and small HDL particles generally decreased by modest amounts, with the exception of triglycerides in these particles which increased 1.5 to 3SD, and total and esterified cholesterol in small HDL which increased by ∼0.1SD. Total, remnant, esterified and free cholesterol, total triglycerides, phosphoglycerides and cholines and total, linoleic, omega-6, MUFA, PUFA and saturated fatty acids all increased by 2-3SD between 16 and 36 weeks, with more mixed and modest changes in fatty acid ratios. VLDL particle size increased by 1SD, HDL size by 0.1SD and LDL particle size decreased by 0.5SD. There were also marked increases in glucose (2SD), pyruvate (2SD) and the inflammatory marker Glycoprotein acetyls (2.5SD). Alanine, glycine, histadine, isoleucine, phenylalanine, creatinine and all of the ketone bodies increased, whereas glutamine, valine, tyrosine and albumin decreased. These patterns of absolute SD metabolic profile change across pregnancy were similar to those seen in the original units (**sTable 3**), and also for rate of change in original units per 4-weeks in the whole cohort with adjustment for randomised arm (**Figure 2**, **sTable 4**).

**Figure 2:**
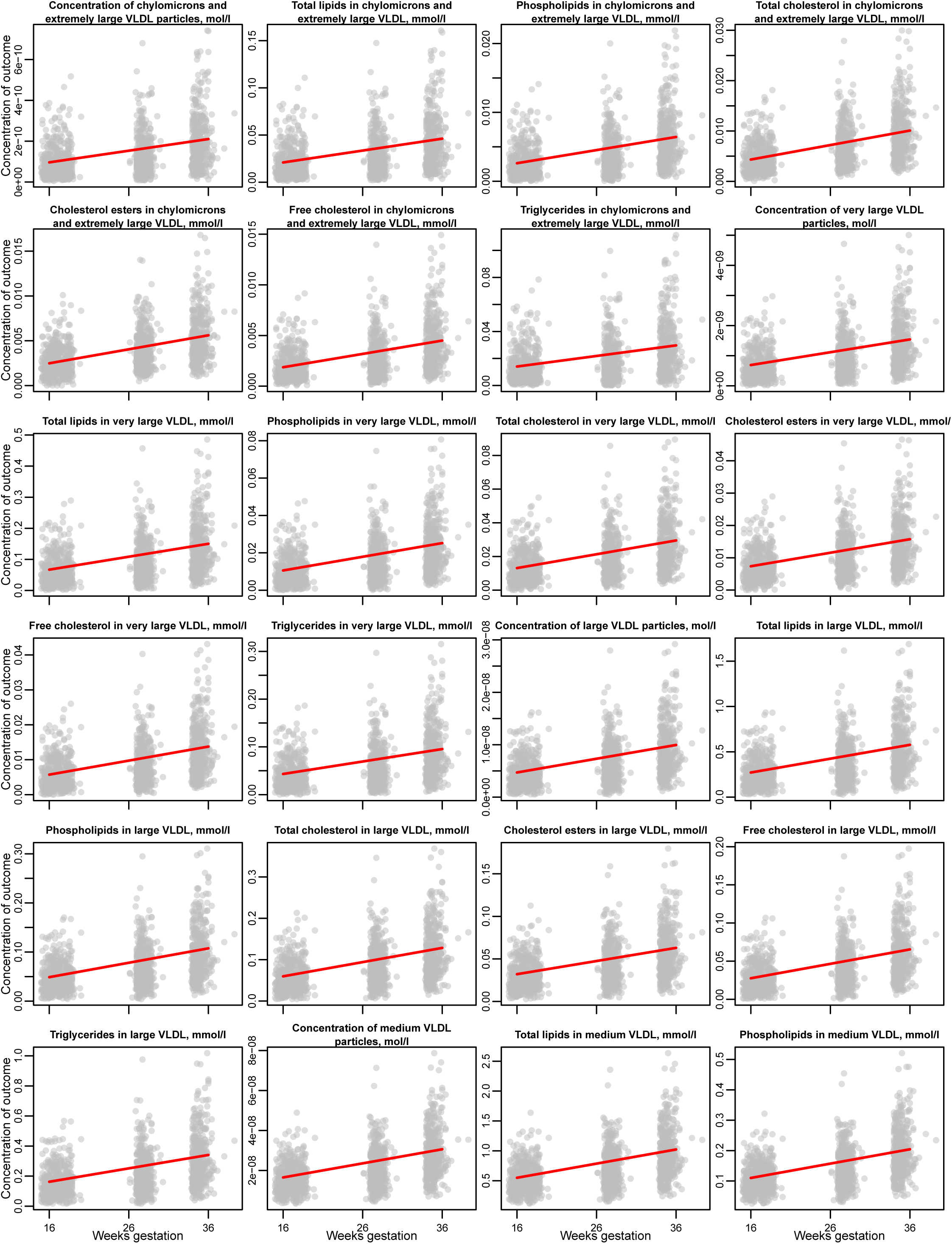
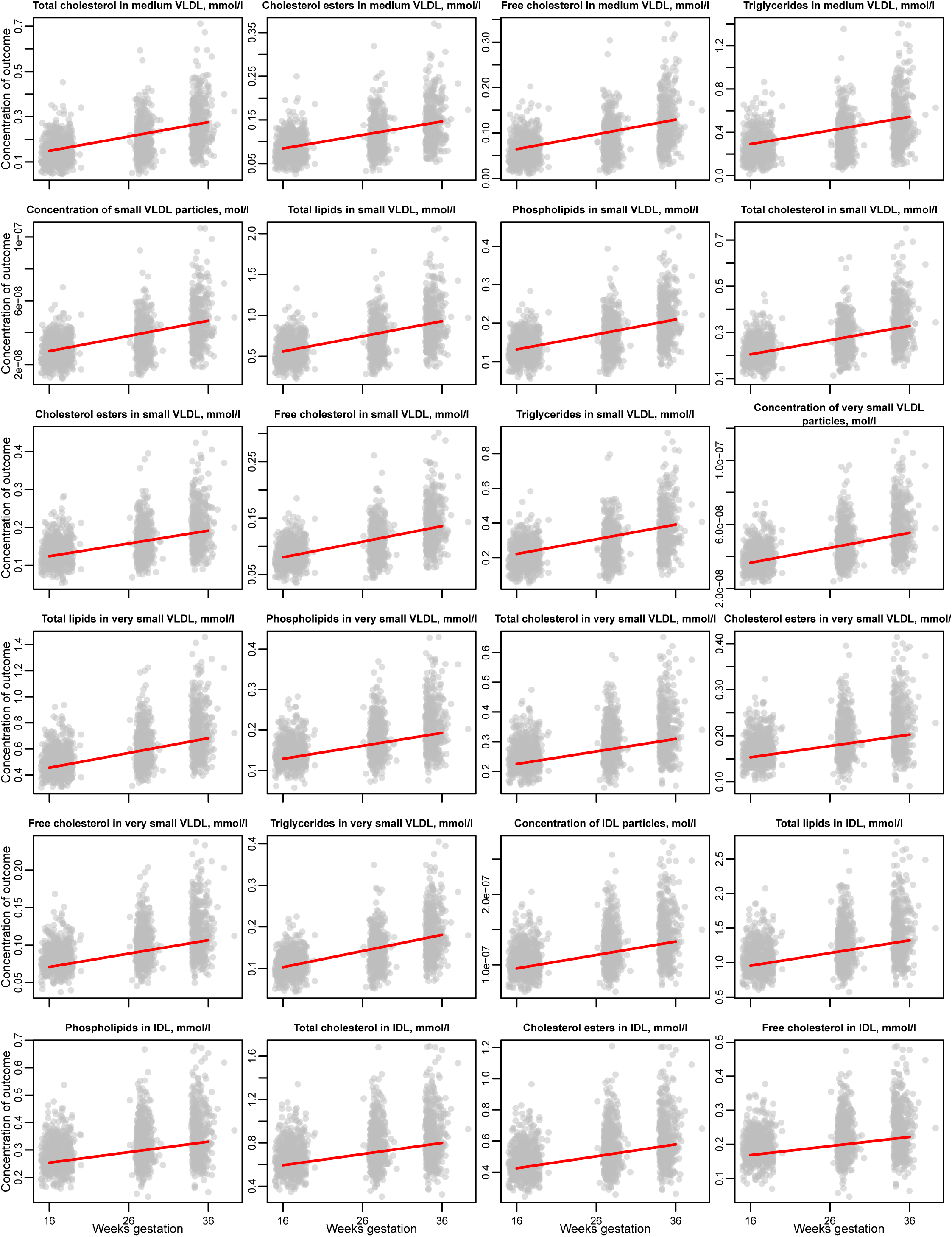
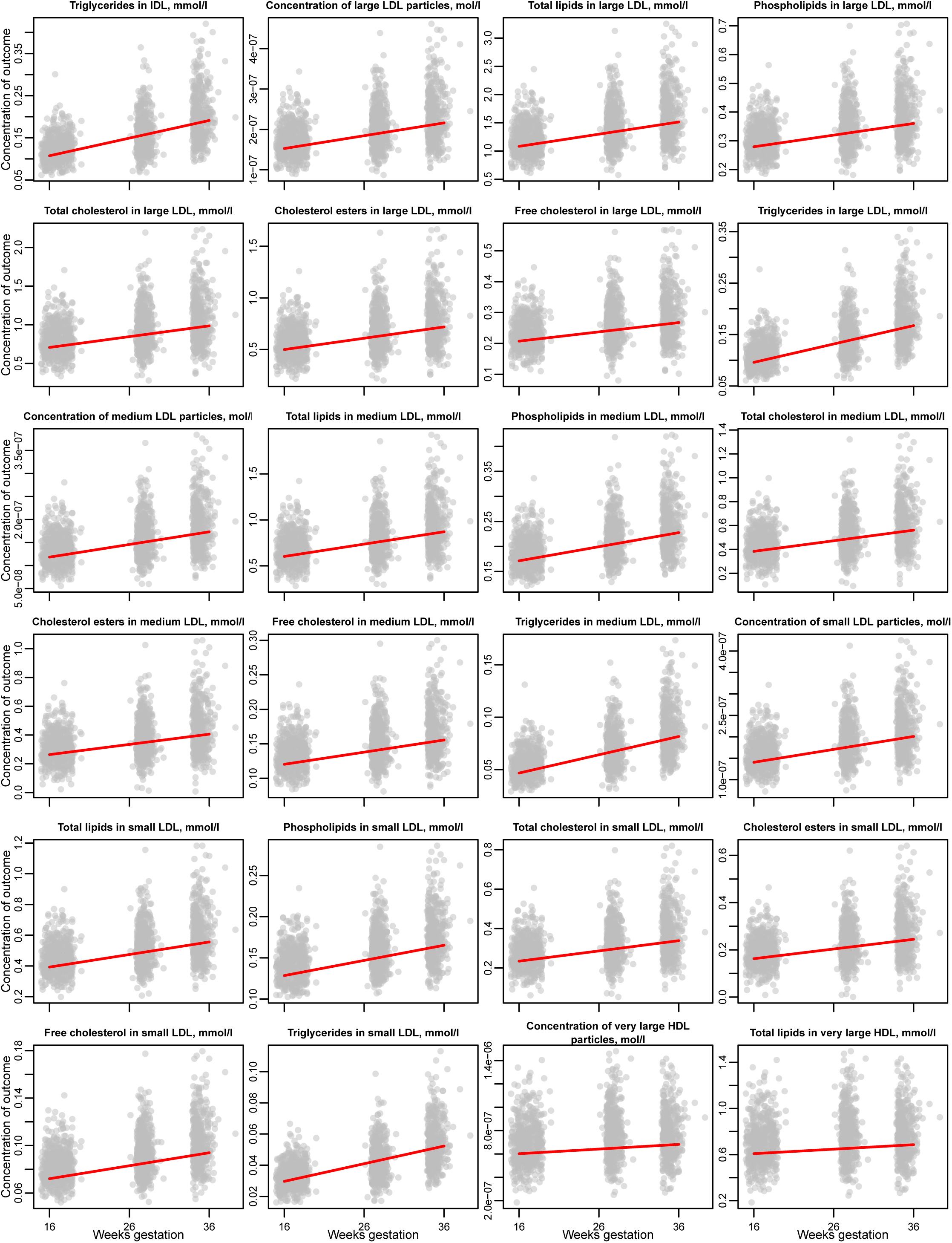
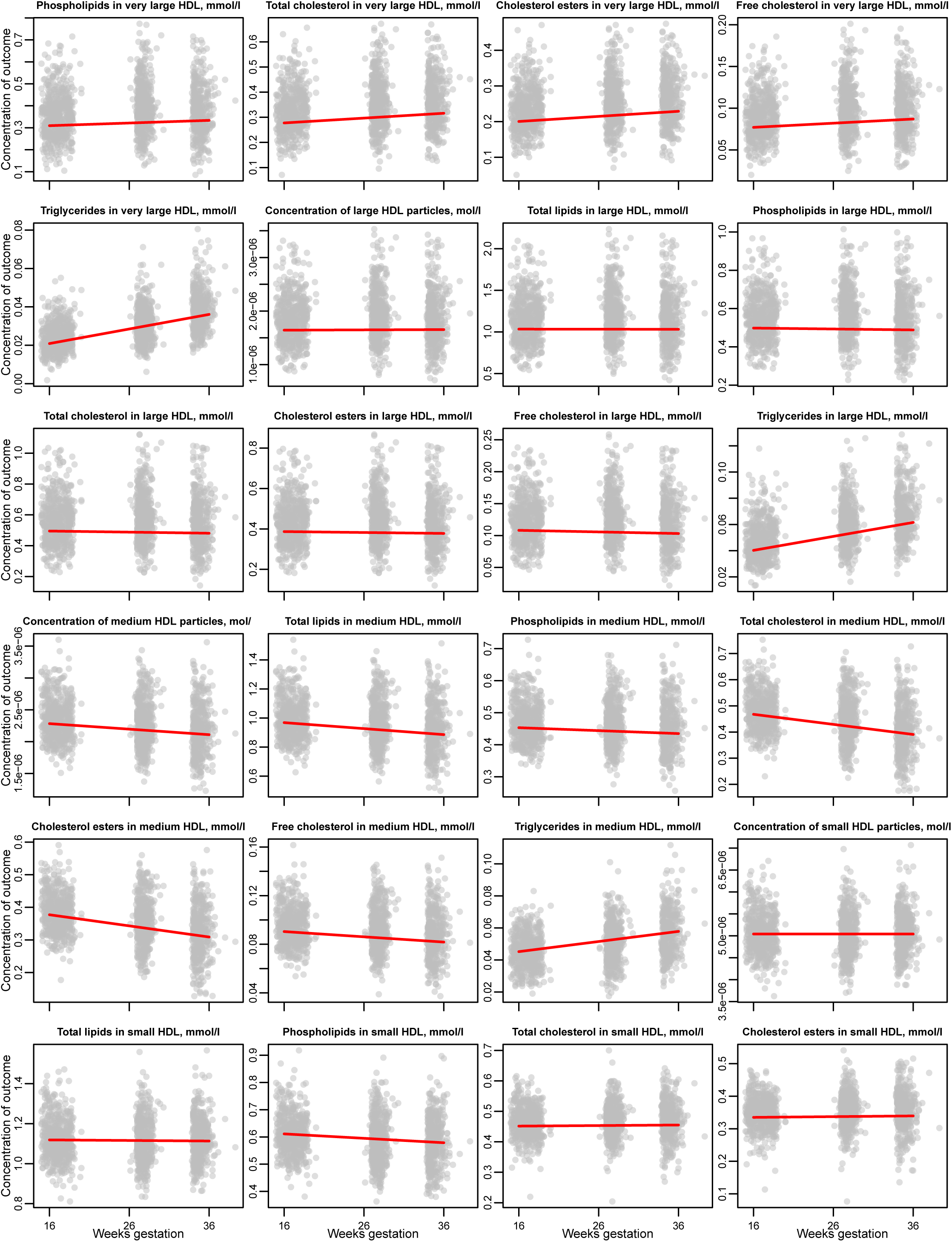
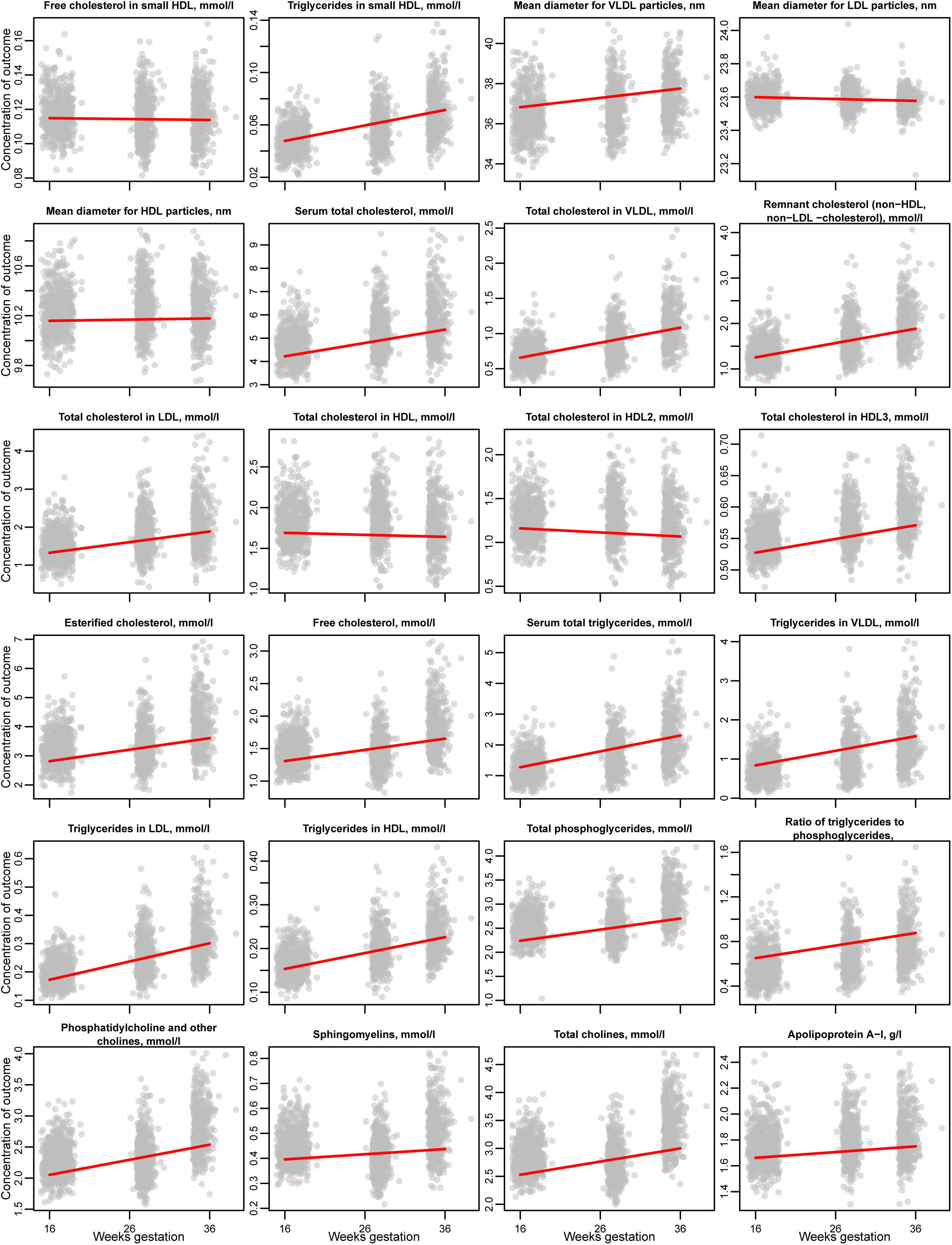
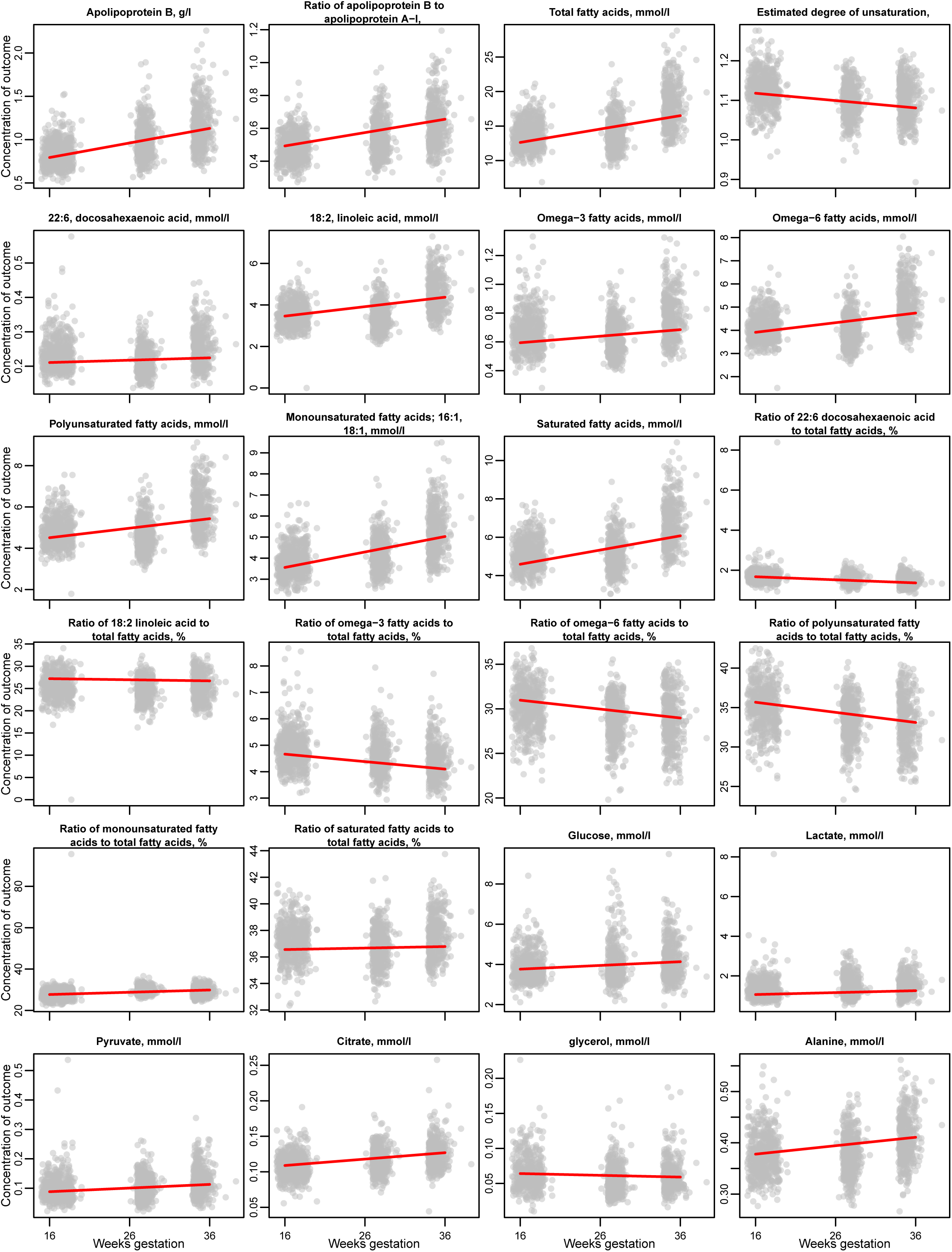
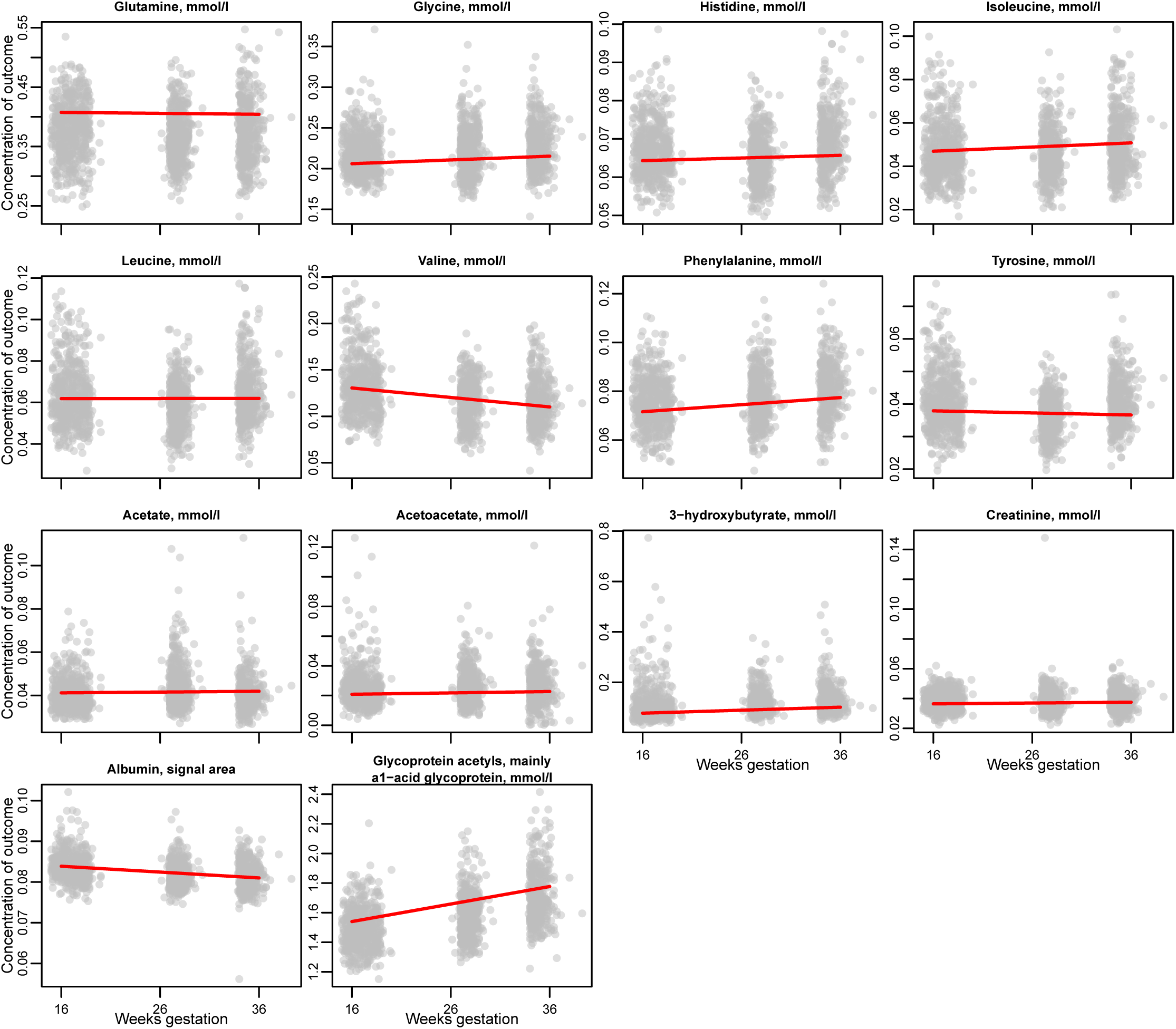
Change in Metabolic trait measurements 16- and 36-weeks of gestation in obese pregnant women (N = 1158). Footnotes to Figure 2: The dots in the figures represent individual measures of each metabolic trait at the three measurement times. The red lines show the fitted predicted rate of change in each trait per 4-weeks of gestation from the multilevel models. These fitted lines are restricted to the time period 16- to 36-weeks. Results are in the original units of each trait. The results are adjusted for any treatment effect by the inclusion of a term for the randomised group in the model. Model predicted numeric results of the absolute change between 16- and 36-weeks of each trait in SD and original units for control women only are shown in Supplementary material sTable 3. The mean value of each trait at 16-weeks and rate of change per 4-weeks between 16- and 36-weeks for all women with adjustment for randomised group are shown in sTable 4.

### Effect of a lifestyle intervention that changed diet and physical activity in obese pregnant women on change in their metabolic profiles

The intervention resulted in reductions in the rate of change of concentrations of all lipids, phospholipids and triglycerides in extremely large, very large, large and medium VLDL particles, except for total cholesterol and cholesterol esters in medium VLDL (**Figure 3, Table 2, sTable 5**). It also resulted in reductions in the rate of change of VLDL particle size, triglycerides in very large HDL, phospholipids in small HDL, and the ratio of total triglycerides to phosphoglycerides. There were effects on fatty acids, with those randomised to the intervention having increased rates of change in the degree of saturation of fatty acids, and the ratios of linoleic, omega-6, and polyunsaturated fatty acids to total fatty acids, and decreased rates of change in the ratio of saturated to total fatty acids. Rates of increase in lactate, pyruvate, and alanine were reduced, and of acetate increased, in those randomised to the intervention (**Figure 3, Table 2, sTable 5**).

**Figure 3:**
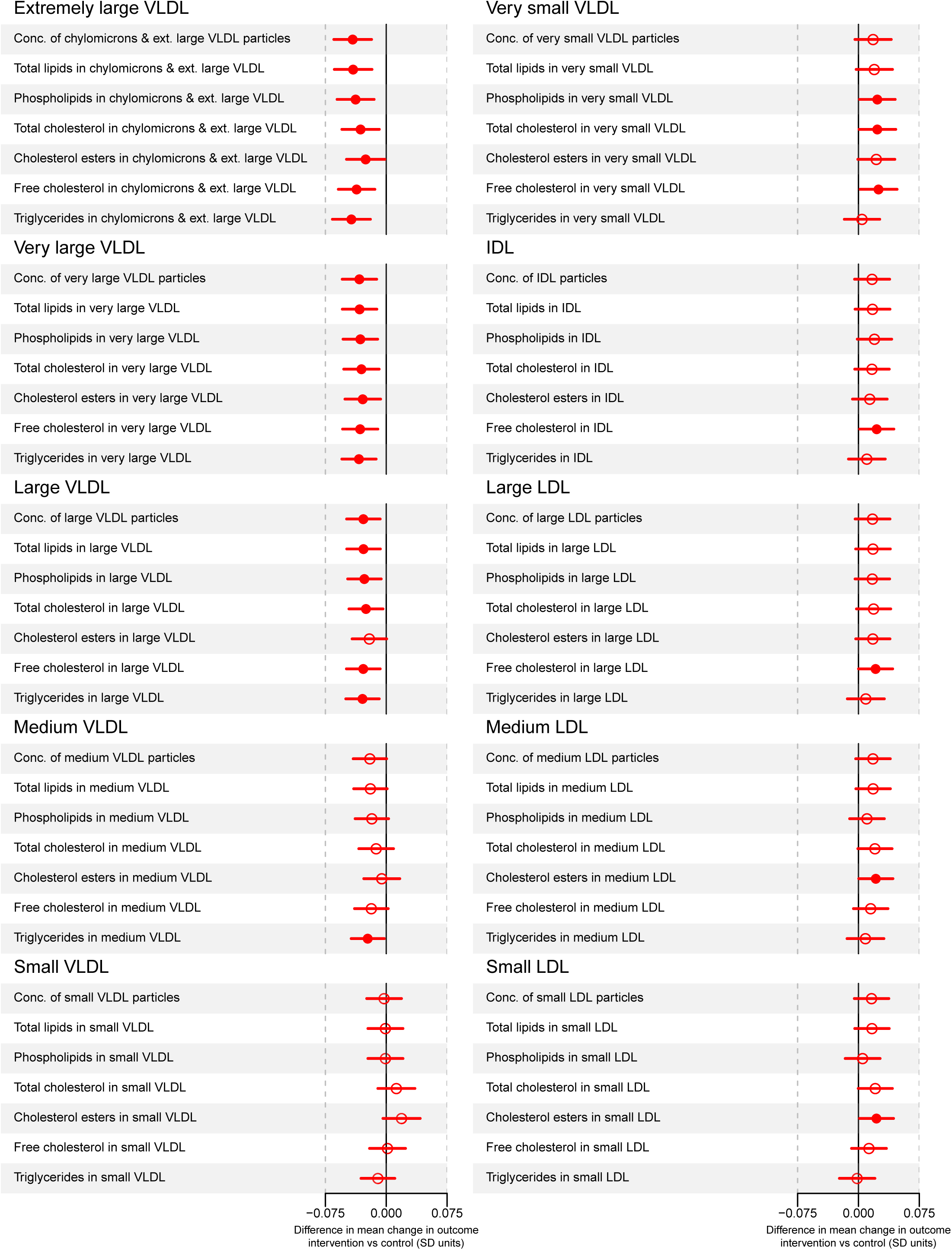
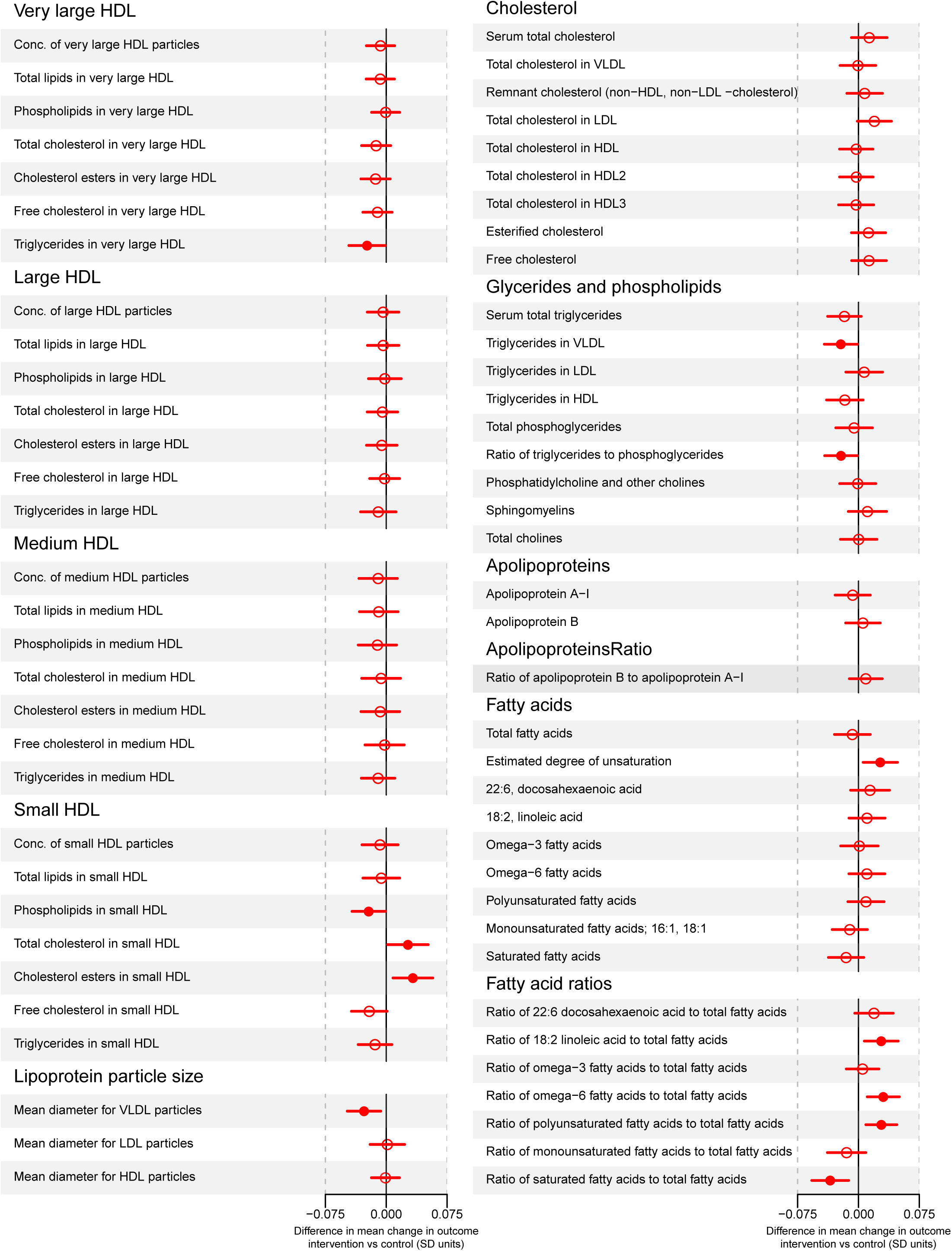
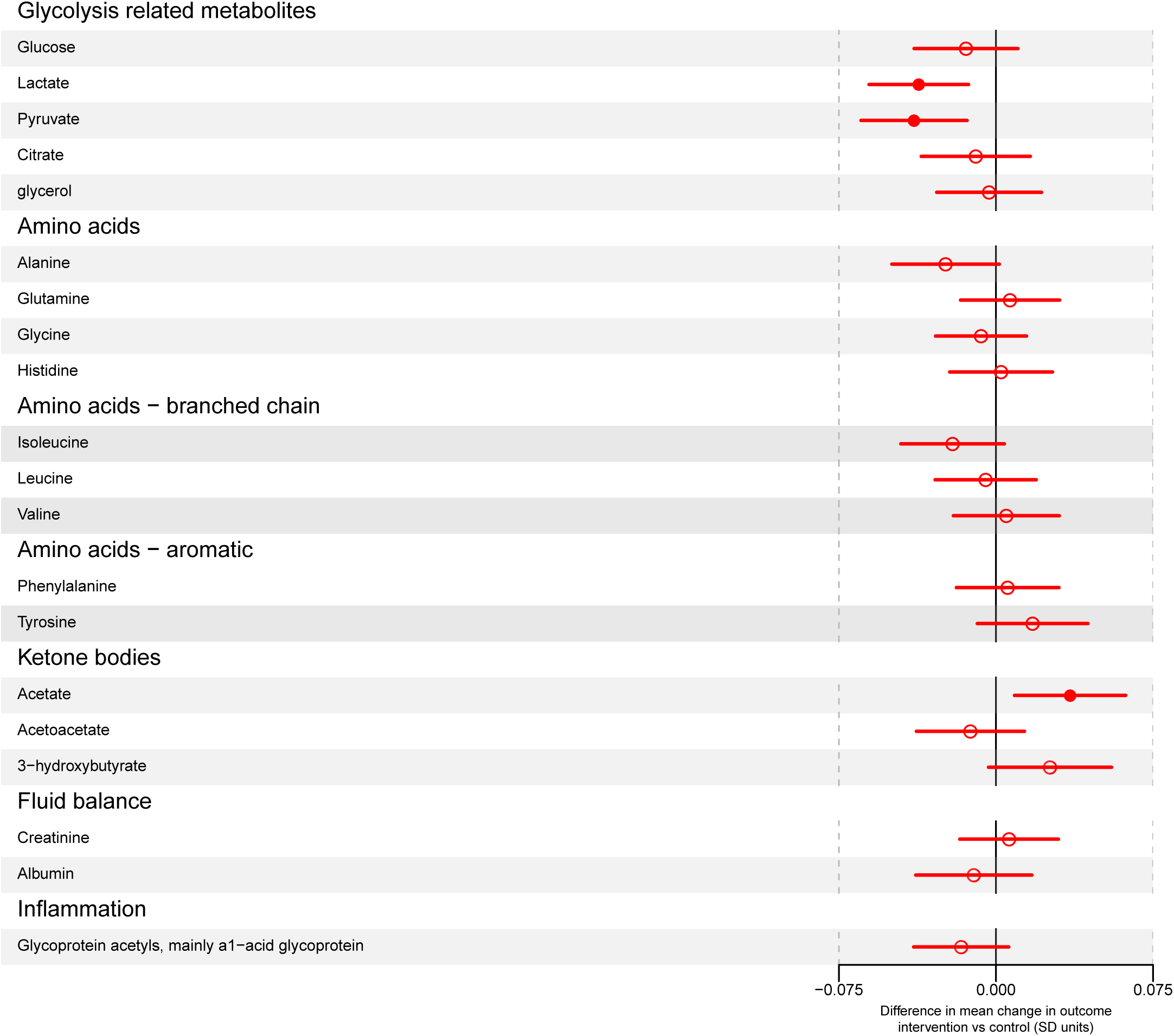
Effect of the UPBEAT intervention on mean rate of change in metabolic traits in SD units per 4-weeks (N = 1158). Footnotes to Figure 3: Circles are the difference in mean rate of change for each metabolic trait in SD units per 4-weeks of gestation comparing those randomised to intervention to those randomised to control (usual antenatal care) and horizontal lines are the 95% confidence intervals for these differences. SDs were calculated from women in the control group. Solid circles are results for which the p-value for difference between the randomised groups is < 0.05; empty circles where it is ≥ 0.05. Results for the differences by randomised group for all traits in the original units (eg. mmol/l) together with absolute p-values are shown in Supplementary material sTable 5.

**Table 2:** Effect of the UPBEAT diet and physical activity lifestyle intervention on selected^a^ metabolic traits. N = 1158.

|  | Difference in mean rate of change in metabolic traits per 4-weeks of gestation between 16 and 36 weeks comparing women receiving intervention to control group |  |
| --- | --- | --- |
|  | *In original units (see first column) per 4-weeks* | *In SD units per 4-weeks<sup>b</sup>* |
| **Extremely large VLDL** |  |  |
| Concentration of chylomicrons and extremely large VLDL particles (mol/l) | -5.430x10<sup>-12</sup><br>(-8.323x10<sup>-12</sup>, -2.537x10<sup>-12</sup>) | -0.041 (-0.065, -0.018) |
| Total lipids in chylomicrons and extremely large VLDL (mmol/l) | -0.001<br>(-0.002, -5.322x10<sup>-4</sup>) | -0.041 (-0.064, -0.018) |
| Phospholipids in chylomicrons and extremely large VLDL (mmol/l) | -1.457x10<sup>-4</sup><br>(-2.287x10<sup>-4</sup>, -6.274x10<sup>-5</sup>) | -0.038 (-0.061, -0.015) |
| Total cholesterol in chylomicrons and extremely large VLDL (mmol/l) | -1.623x10<sup>-4</sup><br>(-2.725x10<sup>-4</sup>, -5.212x10<sup>-5</sup>) | -0.032 (-0.055, -0.009) |
| Cholesterol esters in chylomicrons and extremely large VLDL (mmol/l) | -6.952x10<sup>-5</sup><br>(-1.289x10<sup>-4</sup>, -1.015x10<sup>-5</sup>) | -0.025 (-0.049, -0.002) |
| Free cholesterol in chylomicrons and extremely large VLDL (mmol/l) | -9.276x10<sup>-5</sup><br>(-1.468x10<sup>-4</sup>, -3.868x10<sup>-5</sup>) | -0.037 (-0.059, -0.014) |
| Triglycerides in chylomicrons and extremely large VLDL (mmol/l) | -8.450x10<sup>-4</sup><br>(-0.001, -4.077x10<sup>-4</sup>) | -0.043 (-0.066, -0.020) |
| **Very large VLDL** |  |  |
| Concentration of very large VLDL particles (mol/l) | -2.919x10<sup>-11</sup><br>(-4.656x10<sup>-11</sup>, -1.183x10<sup>-11</sup>) | -0.033 (-0.054, -0.012) |
| Total lipids in very large VLDL (mmol/l) | -0.003 (-0.005, -0.001) | -0.033 (-0.054, -0.012) |
| Phospholipids in very large VLDL (mmol/l) | -4.618x10<sup>-4</sup><br>(-7.453x10<sup>-4</sup>, -1.782x10<sup>-4</sup>) | -0.032 (-0.053, -0.011) |
| Total cholesterol in very large VLDL (mmol/l) | -4.869x10<sup>-4</sup><br>(-8.058x10<sup>-4</sup>, -1.681x10<sup>-4</sup>) | -0.031 (-0.052, -0.009) |
| Cholesterol esters in very large VLDL (mmol/l) | -2.409x10<sup>-4</sup><br>(-4.085x10<sup>-4</sup>, -7.328x10<sup>-5</sup>) | -0.029 (-0.051, -0.007) |
| Free cholesterol in very large VLDL (mmol/l) | -2.459x10<sup>-4</sup><br>(-3.986x10<sup>-4</sup>, -9.316x10<sup>-5</sup>) | -0.032 (-0.054, -0.010) |
| Triglycerides in very large VLDL (mmol/l) | -0.002<br>(-0.003, -7.781x10<sup>-4</sup>) | -0.034 (-0.055, -0.012) |
| **Large VLDL** |  |  |
| Concentration of large VLDL particles (mol/l) | -1.436x10<sup>-10</sup><br>(-2.419x10<sup>-10</sup>, -4.524x10<sup>-11</sup>) | -0.028 (-0.049, -0.008) |
| Total lipids in large VLDL (mmol/l) | -0.008 (-0.014, -0.003) | -0.028 (-0.049, -0.007) |
| Phospholipids in large VLDL (mmol/l) | -0.001 (-0.003, -4.202x10<sup>-4</sup>) | -0.027 (-0.048, -0.006) |
| Total cholesterol in large VLDL (mmol/l) | -0.002 (-0.003, -3.658x10<sup>-4</sup>) | -0.025 (-0.046, -0.004) |
| Cholesterol esters in large VLDL (mmol/l) | -6.322x10<sup>-4</sup><br>(-0.001, -4.684x10<sup>-5</sup>) | -0.021 (-0.042, 5.042x10<sup>-4</sup>) |
| Free cholesterol in large VLDL (mmol/l) | -9.788x10<sup>-4</sup><br>(-0.002, -3.053x10<sup>-4</sup>) | -0.028 (-0.049, -0.007) |
| Triglycerides in large VLDL (mmol/l) | -0.005 (-0.009, -0.002) | -0.029 (-0.050, -0.009) |

**Table 2: continued**
|  |  |  |
| --- | --- | --- |
| **Medium VLDL** |  |  |
| Concentration of medium VLDL particles (mol/l) | -2.706x10<sup>-10</sup><br>(-5.176x10<sup>-10</sup>, -2.363x10<sup>-11</sup>) | -0.020 (-0.041, 4.479x10<sup>-4</sup>) |
| Total lipids in medium VLDL (mmol/l) | -0.009 (-0.017, -5.203x10<sup>-4</sup>) | -0.020 (-0.040, 0.001) |
| Phospholipids in medium VLDL (mmol/l) | -0.002 (-0.003, 2.571x10<sup>-5</sup>) | -0.018 (-0.038, 0.003) |
| Total cholesterol in medium VLDL (mmol/l) | -0.001 (-0.004, 6.072x10<sup>-4</sup>) | -0.012 (-0.034, 0.009) |
| Cholesterol esters in medium VLDL (mmol/l) | -3.934x10<sup>-4</sup><br>(-0.001, 6.747x10<sup>-4</sup>) | -0.006 (-0.028, 0.017) |
| Free cholesterol in medium VLDL (mmol/l) | -0.001 (-0.002, -5.763x10<sup>-6</sup>) | -0.018 (-0.039, 0.002) |
| Triglycerides in medium VLDL (mmol/l) | -0.006 (-0.010, -0.001) | -0.023 (-0.043, -0.002) |
| **Other lipids, lipoproteins affected by the intervention** |  |  |
| Triglycerides in very large HDL (mmol/l) | -2.479x10<sup>-4</sup><br>(-4.714x10<sup>-4</sup>, -2.436x10<sup>-5</sup>) | -0.024 (-0.046, -0.001) |
| Phospholipids in small HDL (mmol/l) | -0.002 (-0.004, -1.363x10<sup>-4</sup>) | -0.022 (-0.042, -0.001) |
| Mean diameter for VLDL particles (nm) | -0.031 (-0.054, -0.008) | -0.027 (-0.048, -0.007) |
| Serum total triglycerides (mol/l) | -0.015 (-0.031, 8.645x10<sup>-4</sup>) | -0.017 (-0.038, 0.003) |
| Triglycerides in VLDL (mmol/l) | -0.015 (-0.028, -0.002) | -0.022 (-0.042, -0.001) |
| Ratio of triglycerides to phosphoglycerides | -0.005 (-0.009, -6.198x10<sup>-4</sup>) | -0.022 (-0.042, -0.001) |
| Estimated degree of unsaturation | 0.001 (1.229x10<sup>-4</sup>, 0.002) | 0.027 (0.006, 0.048) |
| Ratio of 18:2 linoleic acid to total fatty acids (%) | 0.069 (0.013, 0.126) | 0.028 (0.007, 0.049) |
| Ratio of omega-6 fatty acids to total fatty acids (%) | 0.077 (0.023, 0.131) | 0.031 (0.011, 0.051) |
| Ratio of polyunsaturated fatty acids to total fatty acids (%) | 0.079 (0.020, 0.138) | 0.028 (0.009, 0.047) |
| Ratio of saturated fatty acids to total fatty acids (%) | -0.049 (-0.083, -0.015) | -0.035 (-0.058, -0.012) |
| **Other traits affected by the intervention** |  |  |
| Lactate (mmol/l) | -0.017 (-0.027, -0.006) | -0.037 (-0.061, -0.013) |
| Pyruvate (mmol/l) | -0.002<br>(-0.003, -6.028x10<sup>-4</sup>) | -0.039 (-0.064, -0.014) |
| Alanine (mmol/l) | -0.001<br>(-0.002, -2.708x10<sup>-5</sup>) | -0.024 (-0.050, 0.002) |
| Acetate (mmol/l) | 3.081x10<sup>-4</sup><br>(7.541x10<sup>-5</sup>, 5.408x10<sup>-4</sup>) | 0.035 (0.009, 0.062) |

<sup>a</sup>Results are the difference in mean rate of change of each traits in original units (2<sup>nd</sup> column) and SD units (third column) per 4-weeks of gestation between 16 and 36 weeks for selected metabolic traits where p-values are < 0.05. Complete results for all traits assessed in their original units, together with exact p-values, are shown in Supplementary material sTable 5 and in SD units in Figure 3 in the main paper.
<sup>b</sup>SD units are based on the standard deviation values for each metabolite in the control group of women.
SD: standard deviation; VLDL: very low density lipoprotein; LDL: low density lipoprotein; IDL: intermediate density lipoprotein; HDL: high density lipoprotein

### Sensitivity analyses

Though statistical power is reduced in these stratified analyses, the effect of the intervention appeared consistent between 16 and 28 weeks and 28 to 39 weeks (**sFigure 2**).

## Discussion

We have demonstrated marked changes across pregnancy in lipid and glucose metabolic profiles, and in an inflammatory marker, in obese pregnant women. We also found more modest changes across pregnancy in some amino acids, ketone bodies, and metabolites reflecting fluid balance. These findings demonstrate extreme systematic metabolic disruption in obese pregnant women as their pregnancy progresses. Importantly, we have demonstrated that a lifestyle intervention that effectively improved diet and physical activity in these women, resulted in improvements in most VLDL particles and VLDL size. There were also effects on some fatty acid profiles, with increases in the proportion of all fatty acids that were linoleic, omega-6 and polyunsaturated and reductions in the proportion of saturated fatty acids.

An earlier study, which compared the same metabolic profiles as used here between unselected pregnant women and non-pregnant women,^4^ found that lipids and lipoproteins differed by on average 1SD between pregnant and non-pregnant women, with no evidence of a difference in glucose.^4^ In comparison, our findings suggest that the extent of change in lipids, glucose and inflammation across pregnancy in obese pregnant women is more marked than that seen on becoming pregnant. It also appears that the differences in lipids and lipoproteins between women who were in their ^3rd^ and those in their ^2nd^ trimester pregnancies in that previous study were less marked (at most 1SD) than seen in this study of obese pregnant women in this study (up to 3SD). These comparisons support the hypothesis that metabolic profiles are more markedly disrupted in obese than non-obese pregnant women. However, some caution is required in assuming full support for this hypothesis, since our previous study compared pregnant to non-pregnant women and did not have repeat measurements within the same women during pregnancy. To fully quantify differences in metabolic profile changes across pregnancy between obese and non-obese pregnant women requires large prospective studies with repeat assessments of metabolic profiles in both obese and healthy weight women from the same underlying population, and we are not aware of any such studies currently.

The impact of the UPBEAT intervention on VLDLs that we have shown is potentially important for maternal pregnancy adiposity gain and effects on fetal developmental overnutrition.^1^ In normal pregnancy increases in VLDLs, in particular triglycerides in VLDL, result in reduced hepatic lipase activity, the appearance of small dense LDL particles in the maternal circulation and an associated accumulation of maternal hepatic fatty acids. These are a source of fuel for the pregnant woman (ensuring that she deposits sufficient adiposity for healthy fetal development and postnatal breastfeeding), and also an alternative (to glucose) fuel for the fetus.^18^ Thus, substantial increases in VLDLs and triglycerides in obese pregnant women, as observed by us in this study, could result in them gaining more fat during their pregnancy, having a greater propensity for delivery of a LGA infant, and possibly result in greater maternal and offspring adiposity beyond pregnancy and birth. That the intervention tested in this study reduces the increase in VLDLs and triglycerides in obese women might therefore have benefits on risk of maternal and offspring adiposity gain. Whilst this intervention did result in reduced maternal gestational weight gain and adiposity as assessed by skinfolds, supporting our hypothesis of potential benefit for maternal adiposity, it did not influence LGA, which goes against this hypothesis.

Consistent with our finding of no effect of the intervention on rates of increase in glucose levels across pregnancy, the intervention had no impact on GDM.^9^ It is plausible that the lack of effect of this intervention on rate of change in glucose across pregnancy GDM, is explained by the performance of an OGTT at ∼28-weeks, and treatment of those who met criteria for the diagnosis of GDM, in both randomised groups.^9^ This will have similarly reduced glucose after 28 weeks, in those with the highest levels at 28 weeks, in both randomised groups and also reduced the risk of GDM, and GDM associated LGA in both groups.

To our knowledge only one previous RCT has examined the effect of a lifestyle intervention in obese pregnant women on their pregnancy metabolic profiles assessed at 18- to 20-weeks and again at 28- to 30-weeks.^10^ It found no differences in total cholesterol, VLDLc, LDLc, HDLc, triglyceriedes or glucose, at either time point, between those randomised to an intervention aimed at improving physical activity only, one aimed at improving diet or to usual care (control). It did find lower C-reactive protein levels in both intervention groups compared with the control group at 28 to 30-weeks, but not at 18- to 20-weeks.^10^ The considerably smaller sample size and lack of detailed lipid profiles in that previous RCT may explain why they did not find effects on lipid profiles as we have in this study.

### Study strengths and limitations

Key strengths are the application of an intention to treat analysis in a large, well conducted RCT, with concealed random allocation and blinded assessment of metabolic profiles. We have appropriately modelled repeat measurements. These analyses assume that any missing data on the metabolic profiles is missing at random (i.e. that the effect of the intervention in those with some missing metabolic profile data are the same as in those with complete data at all three time points). Given this is a well conducted RCT and there was similar loss to follow-up, and proportions with metabolic profiles at each assessment in the two arms of the trial, it is likely that this assumption is met here. Our main analyses also assumes that changes in metabolites are linear across pregnancy between 16- and 36-weeks. We explored this by comparing change between 16- to 28-weeks to those between 28- to 36-weeks, with results being broadly consistent in these two time periods. However, we acknowledge that statistical power for these two comparisons is limited and they cannot exclude non-linear effects within these periods. Replication of our findings in a similar sized or larger RCT is important to mitigate against these findings being due to chance. However, we are not aware of any other study with relevant data for this replication. The consistency of findings across similar lipoprotein subclasses provides some reassurance that findings are not solely due to chance.

### Conclusions and potential impact

We have shown very marked disruption of metabolic profiles in obese pregnant women, with beneficial effects of a lifestyle intervention that improved their diet and physical activity on VLDL and fatty acid profiles. With further follow-up of these participants we will be able to explore the extent to which these improvements in metabolic profiles impact maternal and offspring adiposity post-natally.

## Acknowledgements

We thank the pregnant women who participated in the UPBEAT study and the UBEAT study team.

## Funding

The UPBEAT trial was funded by the UK’s National Institute for Health Research under its grants for applied research programme (RP-PG-0407-10452). Contributions to funding were also provided by the Chief Scientist Office Scottish Government Health Directorates (Edinburgh) (CZB/A/680), Guys and St Thomas’ Charity (1060508), Tommy’s Charity (SC039280), and the NIHR Biomedical Research Centre at Guy’s and St Thomas’ NHS Foundation Trust and King’s College London.

The research presented in this paper is funded by the European Research Council under the European Union’s Seventh Framework Programme (FP7/2007-2013) ERC grant agreement 669545, the US National Institute of Health (R01 DK10324), the UK Medical Research Council (MRC) (MR/L002477/1) and the NIHR Biomedical Centre at the University Hospitals Bristol NHS Foundation Trust and the University of Bristol. SLW is supported by the Diabetes UK Sir George Alberti Research Training Fellowship. HM, DdSF, KT and DAL work in a Unit that receives support from the UK Medical Research Council (MC_UU_1201/5 and MC_UU_1201/9). LP and DAL are NIHR Senior Investigators (NF-SI-and NF-SI-0166-10196, respectively).

The views expressed are those of the authors and not necessarily those of the UK Medical Research Council, National Health Service, National Institute for Health Research, or Department of Health, or any other funder listed above. The funders had no role in study design, data collection and analysis, decision to publish, or preparation of the manuscript.

## Contributors

All listed authors meet the requirements for authorship. LP, ALB, SMN, NS and PTS contributed to the original UPBEAT trial design. DAL, LP, SMN and NS designed the study that is reported in this paper and obtained funds for collection of metabolite data. DAL, KT, HM and DdSF wrote the analysis plan and HM undertook all main analyses with input from DAL, KT and DdSF; HM and NP completed descriptive analyses for Table 1 and sTable 1. HM, KT and DAL interpreted the results from the statistical analyses. DAL wrote the first draft of the paper. All other authors critically reviewed the first and subsequent drafts. All authors approved the final version, and agree to be accountable for all aspects of the work. DAL is the guarantor of this study.

## Competing interests

Authors declare they have no conflicts of interest.

## Ethics and governance

Approvals were obtained from the UK research ethics committee (UK integrated research application system, reference 09/H0802/5) and local Research and Development (R and D) departments in participating centres; all women provided written informed consent prior to entering the study.

This trial is registered with Current Controlled Trials, ISRCTN89971375.

## References

1. Lawlor DA, Relton C, Sattar N, Nelson SM. Maternal adiposity‐‐a determinant of perinatal and offspring outcomes? Nat Rev Endocrinol 2012; 8(11): 679–88.

2. Liu LX, Arany Z. Maternal cardiac metabolism in pregnancy. Cardiovasc Res 2014; 101(4): 545–53.

3. Lain KY, Catalano PM. Metabolic changes in pregnancy. Clin Obstet Gynecol 2007; 50(4): 938–48.

4. Wang Q, Wurtz P, Auro K, et al. Metabolic profiling of pregnancy: cross-sectional and longitudinal evidence. BMC Med 2016; 14(1): 205.

5. Lawlor DA. The Society for Social Medicine John Pemberton Lecture 2011. Developmental overnutrition‐‐an old hypothesis with new importance? Int J Epidemiol 2013; 42(1): 7–29.

6. Nelson SM, Matthews P, Poston L. Maternal metabolism and obesity: modifiable determinants of pregnancy outcome. Hum ReprodUpdate 2010; 16(3): 255–75.

7. Huda SS, Brodie LE, Sattar N. Obesity in pregnancy: prevalence and metabolic consequences. Semin Fetal Neonatal Med 2010; 15(2): 70–6.

8. Thangaratinam S, Rogozinska E, Jolly K, et al. Effects of interventions in pregnancy on maternal weight and obstetric outcomes: meta-analysis of randomised evidence. BMJ 2012; 344: e2088.

9. Poston L, Bell R, Croker H, et al. Effect of a behavioural intervention in obese pregnant women (the UPBEAT study): a multicentre, randomised controlled trial. The lancet Diabetes & endocrinology 2015; 3(10): 767–77.

10. Renault KM, Carlsen EM, Haedersdal S, et al. Impact of lifestyle intervention for obese women during pregnancy on maternal metabolic and inflammatory markers. Int J Obes (Lond) 2017.

11. Wurtz P, Kangas AJ, Soininen P, Lawlor DA, Davey Smith G, Ala-Korpela M. Quantitative serum NMR metabolomics in large-scale epidemiology: a primer on –omic technology˙. American Journal Epidemiology 2017.

12. Wang Q, Wurtz P, Auro K, et al. Effects of hormonal contraception on systemic metabolism: cross-sectional and longitudinal evidence. Int J Epidemiol 2016; 45(5): 1445–57.

13. Wurtz P, Havulinna AS, Soininen P, et al. Metabolite profiling and cardiovascular event risk: a prospective study of 3 population-based cohorts. Circulation 2015; 131(9): 774–85.

14. Wurtz P, Wang Q, Niironen M, et al. Metabolic signatures of birthweight in 18 288 adolescents and adults. Int J Epidemiol 2016; 45(5): 1539–50.

15. Wurtz P, Wang Q, Soininen P, et al. Metabolomic Profiling of Statin Use and Genetic Inhibition of HMG-CoA Reductase. J Am Coll Cardiol 2016; 67(10): 1200–10.

16. Howe LD, Tilling K, Matijasevich A, et al. Linear spline multilevel models for summarising childhood growth trajectories: A guide to their application using examples from five birth cohorts. Stat Methods Med Res 2016; 25: 1854–74.

17. Tilling K, Macdonald-Wallis C, Lawlor DA, Hughes RA, Howe LD. Modelling childhood growth using fractional polynomials and linear splines. Ann Nutr Metab 2014; 65(2-3): 129–38.

18. Ghio A, Bertolotto A, Resi V, Volpe L, Di Cianni G. Triglyceride metabolism in pregnancy. Adv Clin Chem 2011; 55: 133–53.

